# TORC2 Affects Endocytosis through Plasma Membrane Tension

**DOI:** 10.1101/520551

**Authors:** Margot Riggi, Mariano Macchione, Stefan Matile, Robbie Loewith, Aurélien Roux

**Author notes:** Correspondence to Robbie Loewith or Aurélien Roux.

## Abstract

Target Of Rapamycin complex 2 (TORC2) is a conserved protein kinase that regulates multiple plasma membrane (PM)-related processes including endocytosis. Direct, chemical inhibition of TORC2 arrests endocytosis but with kinetics that are relatively slow and therefore inconsistent with signaling being mediated solely through simple phosphorylation cascades. Here, we show that, in addition to regulation of the phosphorylation of endocytic proteins, TORC2 also controls endocytosis by modulating PM tension. Elevated PM tension, upon TORC2 inhibition, impinges on endocytosis at two different levels: first, by severing the bonds between the PM adaptor proteins Sla2 and Ent1 and the actin cytoskeleton; and, second, by hindering recruitment of Rvs167, an N-BAR-containing protein important for vesicle fission, to endocytosis sites. These results underline the importance of biophysical cues in the regulation of cellular and molecular processes.

## Introduction

Endocytosis is the process by which eukaryotic cells internalize material and information from their environment, and recycle plasma membrane (PM) lipids, trafficking proteins and cell-surface receptors. Membrane remodeling by a well-established sequence of protein complexes (Fig 1) is essential to form endocytic buds that will internalize material (Kaksonen and Roux, 2018). Thus, the PM can be considered as a core part of the endocytic machinery. It is now broadly accepted that physical forces, in particular PM tension, participate in the regulation of the balance between exocytosis and endocytosis in various systems (Dai and Sheetz, 1995) (Gauthier et al., 2012). Functioning in a homeostatic feedback loop, the opposing effects of endocytosis and exocytosis on PM area enable cells to keep tension close to a set point (Apodaca, 2002) (Morris and Homann, 2001) (Fernandez-Sanchez et al., 2015). Additionally, PM tension was shown to regulate specific steps of the endocytosis process, including clathrin pit formation by varying the membrane budding energy (Boulant et al., 2011) (Saleem et al., 2015), and membrane fission by dynamin (Morlot et al., 2012). These tensile forces, depending on the geometry of the bud, constitute either a basal obstacle that the cell machinery has to counteract, or a driving force in order to reshape the PM and form the endocytic vesicle.

**Figure 1:**
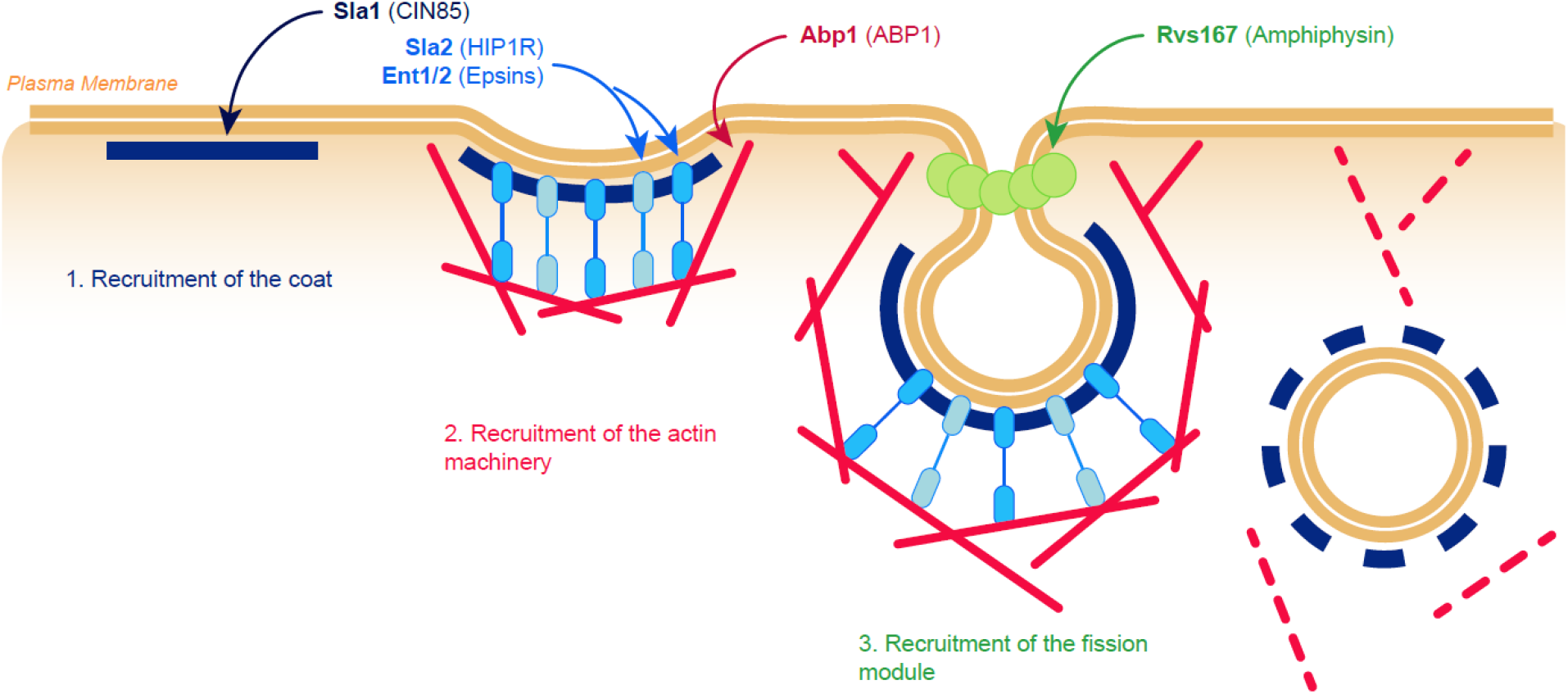
Overview of endocytosis in yeast. Simplified model illustrating the different steps of endocytosis in yeast. The names of the proteins studied in this paper are indicated at the top of the figure. Mammalian orthologs are indicated between brackets.

Membrane remodeling during endocytosis, requires energy. In most mammalian cells, the scaffolding of coat proteins is sufficient to drive membrane invagination. However, in yeast cells which have a high turgor pressure, the dynamic polymerization of actin is additionally required to power PM invagination (Aghamohammadzadeh and Ayscough, 2009; Basu et al., 2014; Dmitrieff and Nedelec, 2015; Kaksonen et al., 2006). In mammalian cells, such an extra force is only needed under conditions when the energy requirements of PM bending are increased, for example at the apical face of polarized epithelial cells (Gottlieb et al., 1993) where membrane bending rigidity is higher, or when membrane tension is increased, as in the case of osmotic swelling or mechanical stretching of cells (Boulant et al., 2011).

The “Target Of Rapamycin Complex 2” (TORC2) was first implicated in the regulation of endocytosis almost 20 years ago when screens for mutants defective in ligand-stimulated internalization of the α-factor receptor identified alleles of *TOR2* and *YPK1*, which encodes an AGC-family kinase and direct substrate of TORC2, (deHart et al., 2002; deHart et al., 2003). The characterization of TORC2 signaling outputs affecting endocytosis was facilitated only recently with the development of chemical-genetic approaches to specifically and acutely inhibit this complex (Gaubitz et al., 2015; Rispal et al., 2015). This revealed that TORC2 regulates endocytosis both through rapid phosphorylation cascades largely mediated by the phospholipid flippase kinases Fpk1 and 2 (Rispal et al., 2015) (Bourgoint et al., 2018; Roelants et al., 2017); and through a slower route presumably involving changes in the biophysical properties of the PM (Rispal et al., 2015).

Independently, we recently demonstrated that TORC2 is a key player in the maintenance of PM tension homeostasis (Berchtold et al., 2012; Riggi et al., 2018), prompting us to speculate that the slower route through which TORC2 regulates endocytosis involves its control of PM tension. Here, we show that this is indeed the case. Inhibition of TORC2 leads to a gradual climb in PM tension, and once a critical threshold is passed, the bonds between the endocytic adaptor proteins and actin filaments fail and coat protein internalization ceases. These phenomena are suppressed if membrane tension is artificially reduced and mimicked if tension is elevated through orthogonal approaches. Lastly, we show that increased tension also blocks the recruitment of Rvs167 to endocytic sites suggesting that the binding of this BAR-domain protein is highly sensitive to membrane curvature.

## Results

### TORC2 inhibition induces the appearance of actin comet tails and the clustering of endocytosis sites

Gradual depletion of TORC2 activity has long been known to lead to an eventual blockade of endocytosis in yeast (deHart et al., 2002; deHart et al., 2003). As endocytosis-related proteins are recruited to, and dissociate from, the budding site in a precise and well-orchestrated order (Kaksonen et al., 2005), we can use them as an internal temporal ladder to study endocytosis dynamics. Here, we followed, as kymographs, the residency times of the actin-binding protein Abp1 at the PM, together with either the coat protein Sla1 or the fission-regulating protein Rvs167 (Fig 2A, B). When we analyzed single endocytic events, we found that in mock-treated cells, consistent with previous results (Kaksonen et al., 2005; Kukulski et al., 2012), all proteins localized to punctate cortical foci. Sla1 resided at an endocytic patch for ~30-40 sec, Abp1 was recruited for a total of ~20 sec, with a ~10 sec overlap with Sla1. Rvs167 was in turn recruited for ~10 sec, and disassembled shortly before Abp1, as the patch budded into the cytoplasm, representing completion of a successful endocytic event (Fig 2C and Supplementary movies 1 and 2). Acute chemical-genetic-inhibition of TORC2 (Bourgoint et al., 2018; Gaubitz et al., 2015) extended all residency times of the proteins at the PM, to the point that endocytic patches often failed to resolve during the time of the experiment (Fig 2A-C and Supplementary movies 3 and 4). Strikingly, we also observed the appearance of Abp1 “comet tails” instead of normal punctate cortical patches. These were anchored to an endocytosis site at the cell cortex - marked by an immobile Sla1 patch – and continuously waved back and forth in the cytoplasm. These structures are typical of an uncoupling between the PM and the actin cytoskeleton (Kaksonen et al., 2003) (Skruzny et al., 2012). Moreover, we observed that most of the blocked endocytic sites were clustered at one given location of the cell, whereas endocytosis events are usually evenly distributed quite homogeneously all along the PM of a (non-budding) cell (Fig 2A, B). Finally, TORC2 inhibition also seemed to affect the recruitment of Rvs167 to the PM, as the corresponding patches were much dimmer and did not always colocalize with Abp1 (Fig 2B, C, Supplementary movie 4). When studying in more detail the kinetics of the endocytosis failure upon TORC2 inhibition (Fig S1), we noticed that we could observe a significant increase in the lifetime of the protein patches only after 30min (Fig 2D-F). The same amount of time was necessary to observe a significant amount of cells displaying clustered endocytosis sites and a “comet tail” phenotype (Fig 2G). Notably, the decrease in intensity of Rvs167 foci was already observable after 15 min, suggesting that it is not a consequence of the blockade of earlier endocytosis steps. These relatively slow kinetics are difficult to explain based solely by changes in protein phosphorylation which occur much faster.

**Figure 2:**
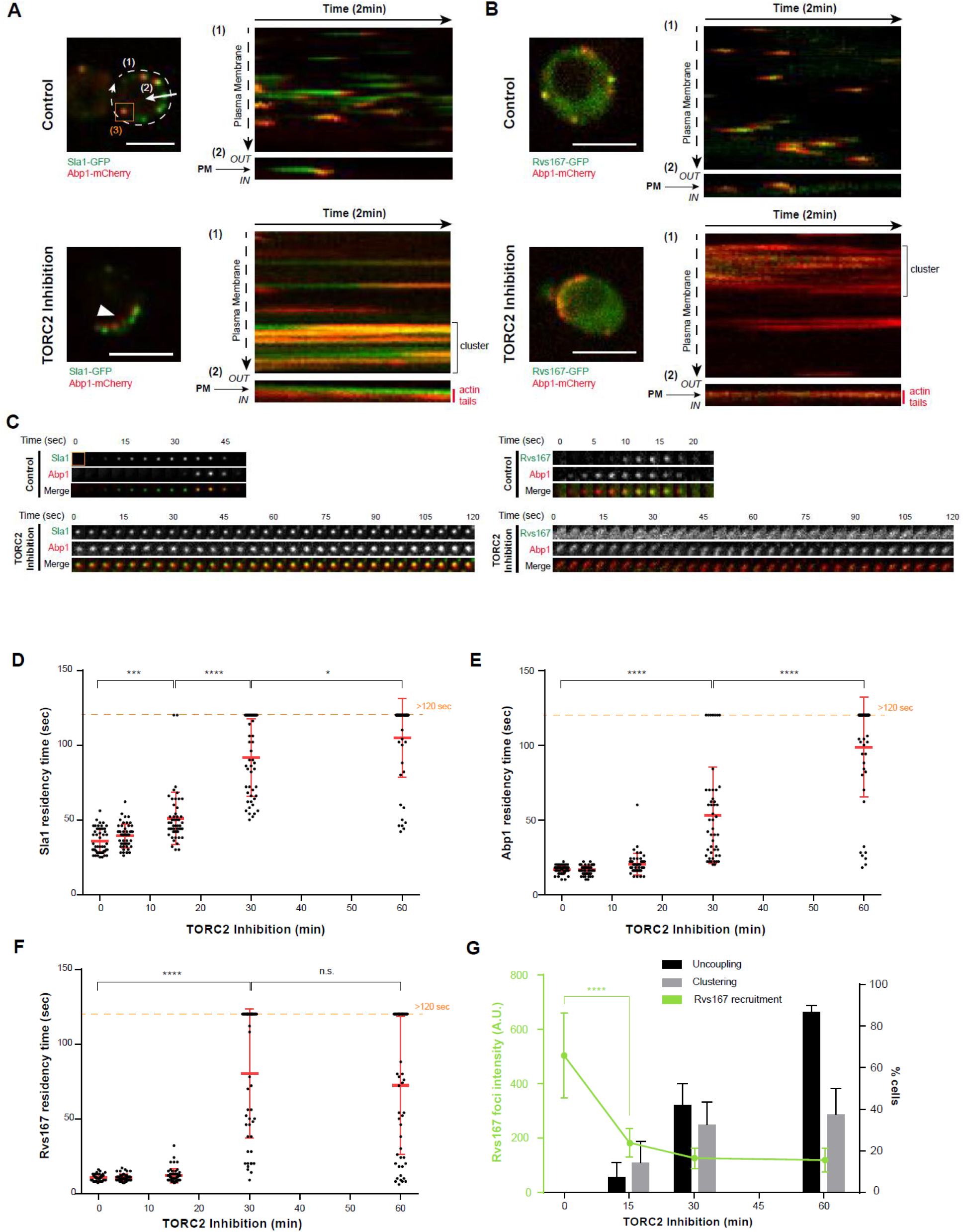
TORC2 inhibition affects endocytosis at multiple stages. **(A, B)** *(Left)* Merged images of Abp1-mCherry and Sla1-GFP **(A)** or Rvs167-GFP **(B)** in *TOR1-1 AVO3Δ^CT^* cells before and after TORC2 inhibition by 1h Rapamycin treatment. *(Right)* Abp1-mCherry and Sla1-GFP (A) or Rvs167-GFP (B) kymographs of the indicated cells, recorded either along (1) or across (2) the Plasma Membrane (PM), as explained on the left pictures. **(C)** Time series of a single endocytic event (as boxed in A) showing Abp1-mCherry signal and either Sla1-GFP (right) or Rvs167-GFP (left), before and after TORC2 inhibition. **(D, E, F)** Evolution of the PM residency times of Sla1 (D), Abp1 (E), and Rvs167 (F), upon TORC2 inhibition in *TOR1-1 AVO3Δ^CT^* cells. Error bars represent s.d. of mean values calculated for n=100 events from at least three independent experiments (**** p<0.0001, * p<0.05). **(G)** Evolution of Rvs167-GFP foci intensity (green plot), and the percentage of cells displaying either an uncoupling phenotype (black boxes) or clustered endocytosis sites (grey boxes) upon TORC2 inhibition in *TOR1-1AVO3Δ^CT^* cells. Error bars represent s.d. of mean values calculated from three independent experiments (with n>100 cells or n>150 foci; **** p<0.0001). All scale bars, 5μm.

### The adaptor proteins Sla2 and Ent1/2 are present at blocked endocytic sites upon TORC2 inhibition

Actin comet tails were previously observed in *SLA2Δ* (Kaksonen et al., 2003) or *ENT1ΔENT2Δ* (Carroll et al., 2012) deletion mutants. These adaptor proteins were shown to cooperatively link the PM to the actin cytoskeleton, efficiently transmitting the force generated by polymerizing actin to the PM. More precisely, Sla2 helps tether Ent1 to the endocytic coat, and, in the absence of Sla2, polymerizing actin pulls Ent1 off the PM (Skruzny et al., 2012). Specifically, the PM remains flat in cells where the Sla2-Ent1/2 link is disrupted (Picco et al., 2018). The binding of Ent1 to actin is regulated through the phosphorylation of five threonines, T346, T364, T395, T415, and T427 (Skruzny et al., 2012) by the Prk1 kinase (Watson et al., 2001). In parallel, both Ent1 and Ent2, as well as Prk1-family proteins, were shown to be hypophosphorylated upon TORC2 inhibition in a phosphoproteomic screen (Rispal et al., 2015; Roelants et al., 2017). This was confirmed in targeted assays in the case of Ent1 (Bourgoint et al., 2018; Rispal et al., 2015).

We thus wondered whether TORC2 inhibition would also affect the behavior of the adaptor proteins Ent1 and Sla2. In untreated cells, the timing of Ent1 recruitment to the PM compared to Sla1 seemed quite variable (Fig 3A), which is consistent with the proposed greater flexibility of the early stages of endocytosis (Godlee and Kaksonen, 2013); in contrast, Ent1 patches assembled on average 25 sec before Abp1 recruitment (Fig 3B). Upon TORC2 inhibition, Ent1 was still present at the PM and localized to blocked endocytic patches, here again often clustered to one position of the cell. It did not travel along the actin comet tails marked with Abp1, but rather remained at the PM with the nonmotile Sla1 patches (Fig 3A-B). Sla2 was similarly progressively blocked at the PM upon TORC2 inhibition, and also did not colocalize with actin comet tails (Fig S2).

**Figure 3:**
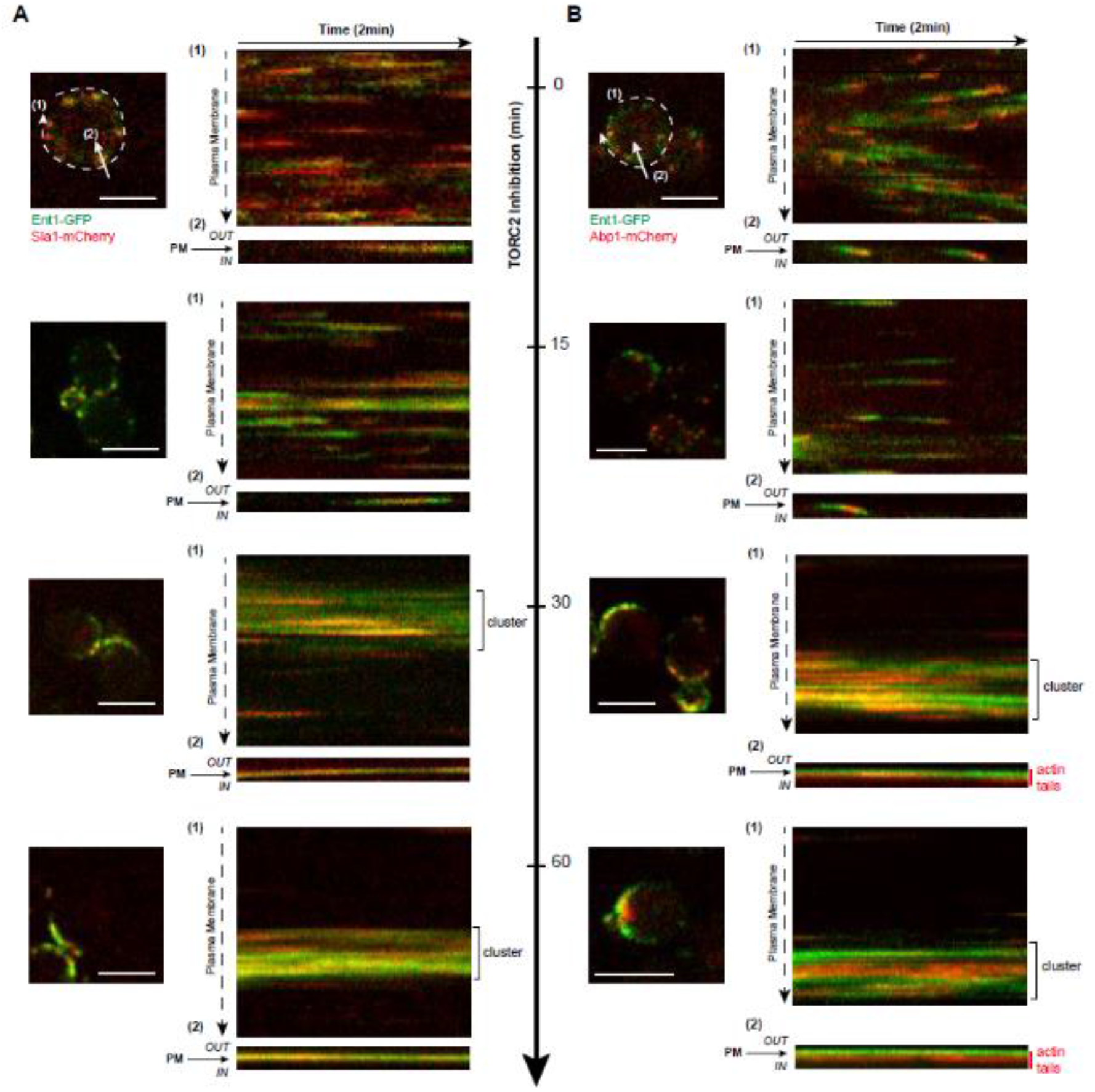
Ent1 colocalizes with blocked endocytosis sites, but not actin tails, upon TORC2 inhibition. **(A, B)** *(Left)* Merged images of Ent1-GFP and Sla1-GFP **(A)** or Abp1-mCherry **(B)** in *TOR1-1 AVO3Δ^CT^* cells upon TORC2 inhibition by Rapamycin treatment. *(Right)* Ent1-GFP and Sla1 - mCherry **(A)** or Abp1-mCherry **(B)** kymographs of the indicated cells, after the indicated time of TORC2 inhibition. Kymographs were recorded either along (1) or across (2) the Plasma Membrane (PM), as explained on the left pictures. All scale bars, 5μm.

These observations suggest that the PM/cytoskeleton uncoupling phenotype linked to TORC2 inhibition is due to defects in the link between the adaptor proteins and the actin cytoskeleton, and not between the adaptor proteins and the PM.

### The actin comet tail phenotype associated with TORC2 inhibition is due to increased PM tension

The slow and progressive appearance of clustered actin comet tails amongst a population of cells suggests that it is unlikely only due to an inhibition of discrete phosphorylation cascade downstream of TORC2. We rather hypothesized that it is the consequence of the increase of PM tension that follows TORC2 inhibition (Riggi et al., 2018). This parameter can now be easily evaluated by Fluorescent Lifetime Imaging Microscopy (FLIM) through the lifetime of the Flipper-TR probe (Colom et al., 2018) and we observed that both phenomena (appearance of actin comet tails and rise in PM tension) follow a similar kinetics (Fig 4A).

**Figure 4:**
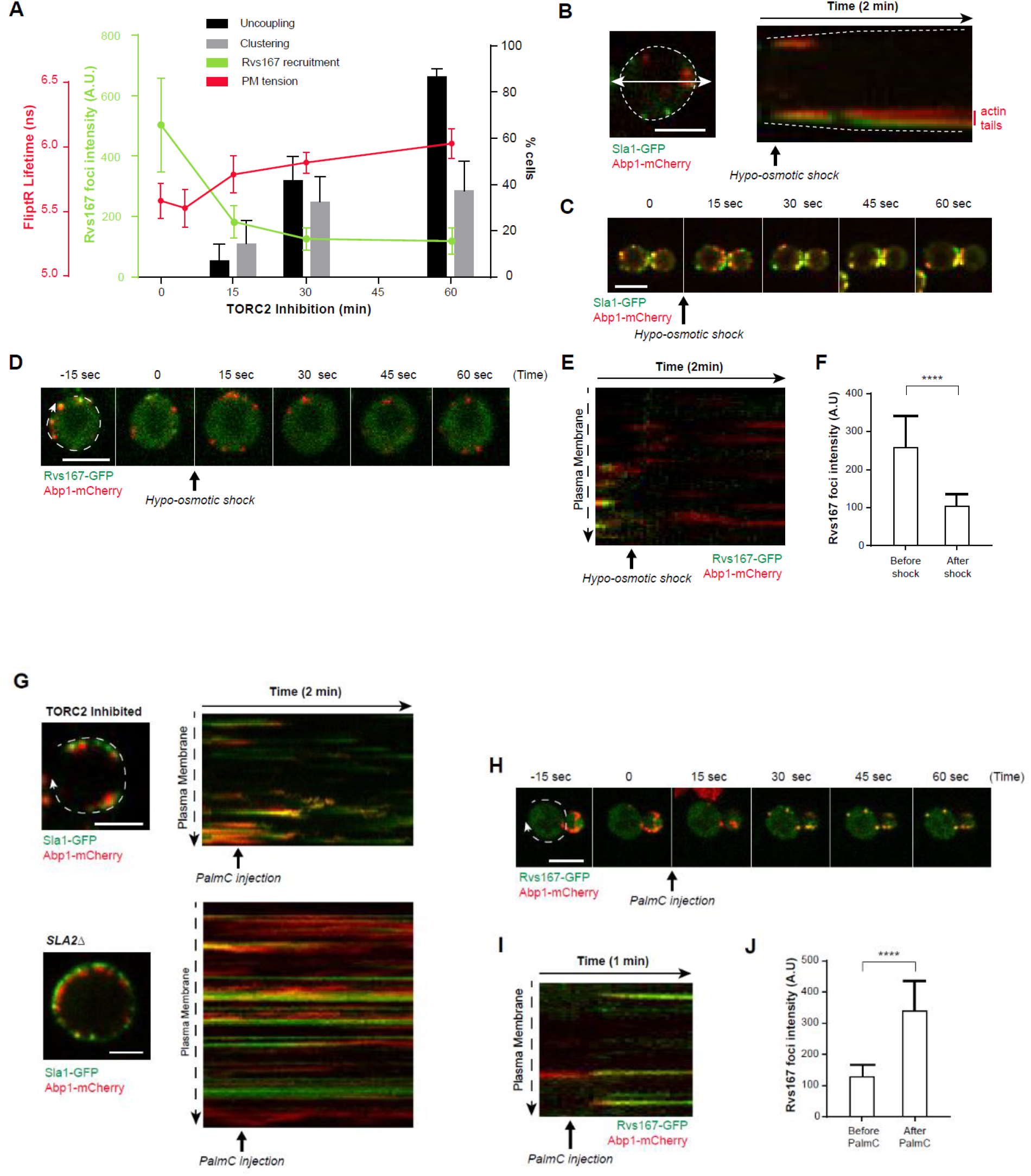
The uncoupling phenotype and Rvs167 misrecruitment are due to an increase in Plasma Membrane (PM) tension. **(A)** Correlation between the kinetics of the increase in PM tension upon TORC2 inhibition, and of the appearance of the various endocytosis phenotypes. PM tension was monitored through the lifetime of the FliptR probe measured by FLIM, and error bars represent the propagated error of mean values calculated from three independent experiments (with n>10 cells). **(B)** Sla1-GFP and Abp1-mCherry kymograph recorded as indicated across a FPS1 cell upon hypo-osmotic shock. Dot lines represent the PM. **(C)** Time series of merged Sla1-GFP and Abp1-mCherry signals in FPS1 cells upon hypo-osmotic shock. **(D)** Time series of merged Rvs167-GFP and Abp1-mCherry signals in a *FPS1Δ* cell upon hypo-osmotic shock. **(E)** Rvs167-GFP and Abp1-mCherry kymograph recorded as indicated in (D) along the PM of a *FPS1Δ* cell upon hypo-osmotic shock. **(F)** Average intensity of Rvs167-GFP foci before and after hypo-osmotic shock. Error bars represent s.d. of mean values calculated for n=100 events from three independent experiments (**** p<0.0001). **(G)** Sla1-GFP and Abp1-mCherry kymographs recorded as indicated along the PM of a *TOR1-1 AVO3Δ^CT^* cell, deleted (bottom) or not (top) of *SLA2*, and pre-treated with Rapamycin for 1h, upon PalmC injection. **(H)** Time series of merged Rvs167-GFP and Abp1-mCherry signals upon PalmC injection in a *TOR1-1 AVO3Δ^CT^* cell, pre-treated with Rapamycin for 1h. **(I)** Rvs167-GFP and Abp1-mCherry kymograph recorded as indicated in (H) along the PM of a *TOR1-1 AVO3Δ^CT^* cell, pre-treated with Rapamycin for 1h, upon PalmC injection. **(J)** Average intensity of Rvs167-GFP foci before and after PalmC treatment in a *TOR1-1 AVO3Δ^CT^* cell, pre-treated with Rapamycin for 1h. Error bars represent s.d. of mean values calculated for n=100 events from three independent experiments (**** p<0.0001). All scale bars, 5μm.

To test this assumption, we investigated whether increased PM tension induced through orthologous means could trigger a similar phenotype. We recently demonstrated that osmotic shocks can be used to manipulate PM tension in yeast (Riggi et al., 2018). As *WT* yeast cells adapt within a few minutes to a hypo-osmotic shock, we used *FPS1Δ* cells that cannot efficiently export the osmo-protectant glycerol (Beese et al., 2009) – indeed, upon hypo-osmotic shocks their PM tension does not increase significantly more than in *WT* cells, but is slower to recover (Fig S3A). As hypothesized, a hypo-osmotic shock induced the formation of actin comet tails anchored to a blocked endocytic site marked by a non-motile Sla1 patch (Fig 4B and supplementary movie 5), and the clustering of endocytosis sites (Fig 4C).

A hypo-osmotic shock also mimicked the defects in Rvs167 recruitment at the PM (Fig 4D, E and supplementary movie 6), and the foci that still formed after the shock were on average half as bright (Fig 4F). These phenomena were similar to those observed upon TORC2 inhibition but differed in one important regard: they were manifest within seconds of treatment (in contrast to the 15-30 minutes required following TORC2 inhibition).

Conversely, we examined whether the uncoupling phenotype caused by TORC2 inhibition could be rescued by artificially decreasing PM tension through Palmitoylcarnitine (PalmC) treatment (Riggi et al., 2018). Although the lifetime of the endocytic patches at the PM remained on average longer than in control cells - likely because TORC2 also regulates endocytosis through other pathways (i.e. phosphorylation cascades) which are still affected - PalmC addition triggered the restart of all frozen endocytic events, and induced the disassembly of the actin comet tails and the redistribution of clustered endocytic patches (Fig 4G and supplementary movie 7). This suggests that these latter two features are controlled by TORC2 *via* PM tension – whereas the elongation of the patch lifetime is likely also influenced by the downregulation of related phosphorylation cascades.

In accordance with a previous study which could rescue endocytosis defects with hyper-osmotic shocks in various mutants, but not in *SLA2Δ* cells, (Basu et al., 2014), PalmC treatment did not rescue the uncoupling phenotype due to *SLA2* deletion (Fig 4G). As *SLA2Δ* cells do not exhibit a particularly high PM tension under basal conditions and still exhibit a decrease of PM tension upon PalmC treatment (Fig S3B), we hypothesize that, in this case, the phenotype is rather due to the complete loss of the physical link between the PM and the cytoskeleton. Consistently, Ent1 has been shown to require Sla2 to stably bind to the PM (Skruzny et al., 2012; Skruzny et al., 2015).

Decreasing PM tension through PalmC treatment could also partially rescue the defects in Rvs167 recruitment (Fig 4H and Supplementary movie 8); even if the residency time of the patches was still longer than in basal conditions (Fig 4I), they became, within a few seconds, more than twice brighter (Fig 4I, J).

Together, these observations demonstrate that both PM tension increase (caused by TORC2 inhibition) and the loss of the connecting bridge between the PM and the actin cytoskeleton, trigger PM/cytoskeleton uncoupling. Moreover, they confirm the importance of the level of PM tension for an efficient recruitment of Rvs167.

### Actin tails are the consequence of an imbalance between PM tension and the strength of its link to the cytoskeleton

We hypothesized that if PM tension becomes too high then the bonds between the PM and adaptor proteins and/or adaptor proteins and actin filaments might be unable to support the force necessary to invaginate the membrane. To test this idea, we wondered whether weakening the link that couples the PM and the cytoskeleton would result in a faster appearance of the actin tails upon TORC2 inhibition. Single deletions of *ENT1* or *ENT2*, which encode epsin homologs, does not cause any obvious phenotype (Wendland et al., 1999), however it might be enough to destabilize the connection between the PM and the cytoskeleton. When we followed the evolution of PM tension upon TORC2 inhibition in *ENT1Δ* cells, we could not detect any significant difference compared to *WT* cells (Fig 5A). However, the appearance of the extension of the PM residency time of Sla1 (Fig 5B) and Abp1 (Fig 5C), as well as the appearance of the uncoupling and the clustering phenotypes were all faster (Fig 5D), meaning that a lower PM tension is sufficient to break the remaining bonds between the PM and the cytoskeleton.

**Figure 5:**
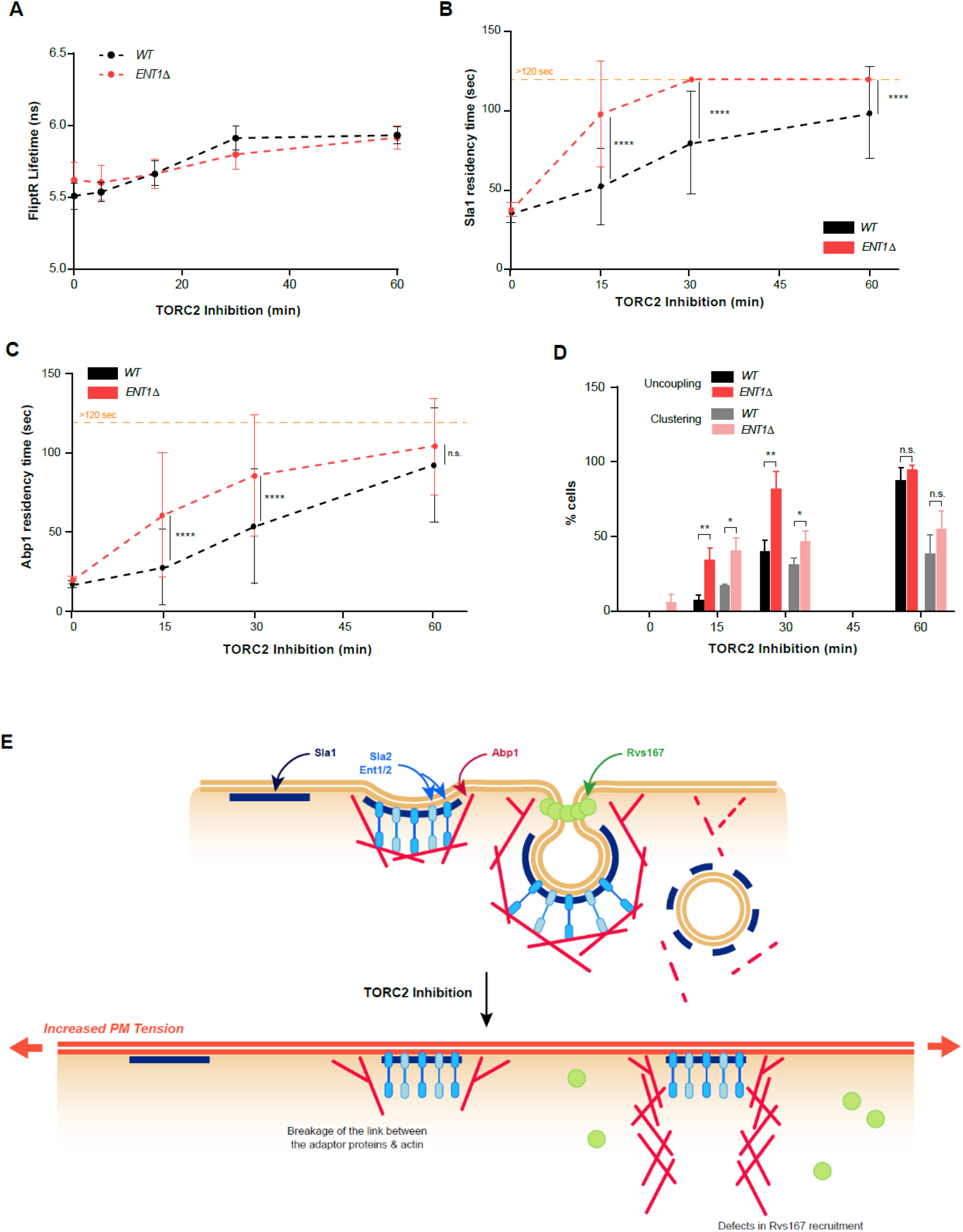
The uncoupling phenotype is the consequence of an unbalance between Plasma Membrane (PM) tension and the force which the adaptor proteins can sustain. **(A)** Evolution of PM tension, monitored through the lifetime of the FliptR probe measured by FLIM, upon TORC2 inhibition in *TOR1-1 AVO3Δ*^CT^ cells deleted or not of *ENT1*. Error bars represent the propagated error of mean values calculated from three independent experiments (with n>10 cells). **(B, C)** Comparison of the evolution of the PM residency times of Sla1 **(B)**, or Abp1 **(C)**, upon TORC2 inhibition in *TOR1-1 AVO3Δ*^CT^ cells deleted or not of *ENT1*. Error bars represent s.d. of mean values calculated for n=100 events from at least three independent experiments (**** p<0.0001). **(D)** Comparison of the kinetics of appearance of the uncoupling phenotype upon TORC2 inhibition in *TOR1-1 AVO3Δ*^CT^ cells deleted or not of *ENT1*. Error bars represent s.d. of mean values calculated for n=100 events from at least three independent experiments (* p<0.05, ** p<0.01). (E) Model illustrating that TORC2 inhibition induces an increase in PM tension that affects endocytosis at several levels.

Together, these results that PM tension is a crucial parameter governing endocytosis in yeast. Indeed, the connecting bridge transmitting the force developed by the cytoskeleton to actively reshape the PM and pull it inwards can only resist a defined threshold of tension, above which it breaks and actin polymerization becomes futile.

## Discussion

Previous work had already demonstrated that TORC2 regulates endocytosis both through direct phosphorylation cascades largely mediated by the phospholipid flippase kinases Fpk1 and 2; and through changes in the biophysical properties of the PM (Rispal et al., 2015). Here, we confirmed that some of the endocytic defects observed in TORC2-inhibited cells, specifically the recruitment of Rvs167, and the uncoupling between the PM and the cytoskeleton, are due to an increase in PM tension. More precisely, the latter phenomenon is the consequence of the breakage of the connecting bonds between the adaptor proteins and actin filaments upon increased tension (Fig 5E). This underlines the importance of biophysical cues in the regulation of cellular and molecular processes. Notably, the defect in Rvs167 recruitment to endocytic foci appears faster than all the other phenotypes, whereas the kinetics of the increase of its residency time is similar to other proteins. This suggests that TORC2 might impinge on this process through an additional pathway in parallel to the increase in PM tension. Indeed, Rvs167 binding to the PM is known to be sensitive to PM biophysical properties such as curvature (Simunovic et al., 2015; Youn et al., 2010), but also to its composition and specifically to the level of complex sphingolipids (Youn et al., 2010), which is also regulated by TORC2 (Aronova et al., 2008; Breslow et al., 2010; Muir et al., 2014; Roelants et al., 2011).

We observed that the PM tension has to reach a threshold, and maintain it for an amount of time, before the actin cytoskeleton uncouples from the PM. This threshold can be seen as the maximum force that the link between the adaptor proteins and the cytoskeleton can withstand. This threshold can be artificially decreased by weakening this connection, for example through *ENT1* single deletion. The adaptors Sla2 and Ent1/2 constitute a physical bridge between the PM and the actin cytoskeleton, each of these proteins possessing a PtdIns(4,5)P2-binding domain on their N-terminal part, and in the case of Sla2, a THATCH/talin-like domain at its C-terminal end (Skruzny et al., 2012). The Ent1-actin interaction domain is additionally known to be phosphoregulated by Prk1 (Skruzny et al., 2012), which is also a potential downstream target of TORC2 (Bourgoint et al., 2018; Rispal et al., 2015). However, as the uncoupling phenotype is largely rescued by an artificial decrease of PM tension induced by PalmC – a condition in which TORC2 inhibition is maintained – it is more probably the direct mechanical effect of increased PM tension, rather than, or in addition to, a consequence of the TORC2-related downregulated phosphorylation cascade. Interestingly, it seems that the biophysical cue of increased PM tension can be transmitted through the adaptor proteins, as it affects rather their binding to the actin cytoskeleton than to the PM.

Another striking phenotype associated with TORC2 inhibition is the clustering of the endocytosis sites. It could also be recapitulated by increasing PM tension through orthologous means, and conversely rescued by decreasing PM tension, suggesting that here again increased PM tension is the direct cause of the clustering. We can hypothesize that the assembly of a first endocytosis patch facilitates the assembly of more patches in its vicinity by modifying the biophysical properties of the PM locally. Together, these observations suggest that the uncoupling phenotype can have (at least) two different causes: the deletion of the genes encoding the proteins that connect the PM to the cytoskeleton, or an increase in PM tension; only the latter also induces the clustering of the endocytosis patches. The force developed by actin polymerization, and its efficient transmission to the PM, is of particular importance to power endocytosis in yeast, as the cells have to counteract a high turgor pressure due to their cell wall. Importantly, HeLa cells in which Sla2 mammalian homolog Hip1R is down-regulated display the same actin comet tail phenotype (Engqvist-Goldstein et al., 2004), as well as the cellular slime mold *Dictostelium discoideum* lacking epsins (Brady et al., 2010), meaning that the molecular mechanism of PM-actin cytoskeleton coupling is likely conserved across evolution. It would be interesting to investigate whether this connection is also under the influence of the PM tension in higher organisms, and potentially of TORC2, as it has also been linked to PM tension regulation in higher eukaryotes (Diz-Munoz et al., 2016).

## Material & methods

### Yeast strains and plasmids

All strains used in this study are listed in Supplementary Table 1. Yeast strains were generated either by homologous recombination of PCR-generated fragments as previously described or by crossing, sporulation and subsequent dissection of the spores. All plasmids and primers used for the generation of the strains are listed in Supplementary Tables 2 and 3. Strains were confirmed by PCR and sequencing. Cloning and site-directed mutagenesis were performed following standard procedures and plasmids were verified by sequencing. All tagged proteins are functional and expressed from their endogenous promoter.

### Yeast culture

Yeast cells were grown according to standard procedures at 30°C Synthetic Complete medium to an early logarithmic phase. For the hypo-osmotic shocks, cells were grown in Synthetic Complete medium containing 1 M of sorbitol, and pre-warmed Synthetic Complete medium was injected inside a flow chamber.

### Chemicals

Rapamycin (LC Laboratories) was dissolved in DMSO at 1mg/mL and used at a final concentration of 200nM. Palmitoylcarnitine was dissolved in DMSO at 10mM and used at 5μM. The Flipper-TR probe, synthetized following reported procedures (Colom et al., 2018), was dissolved in DMSO, and used at a final concentration of 2ng/mL.

### Confocal Microscopy

Cells were grown at 30°C in SC medium to an early logarithmic phase, mounted on coverslips coated with 0.1 μg/mL Concanavalin A (Sigma) and immediately imaged with a spinning-disc microscope assembled by 3i (Intelligent Imaging Innovation, Denver, USA) and Nikon (Eclipse C1, Nikon, Tokyo, Japan) using a 100 x objective (NA=1.3, Nikon). For microfluidics experiments, a Concanavalin A-coated coverslip was bonded to the bottom surface of a flow chamber (sticky-slide VI 0.4, Ibidi, Munich, Germany) with one entry connected to a syringe pump (Aladdin, World Precision Instrument, Sarasota, USA) and the other left open for sequential introduction of different solutions. The flow chamber was primed with SC medium prior to the loading of cells. Loaded cells were washed several times with SC medium, and then subjected to the appropriate treatments.

### Fluorescence Lifetime Imaging Microscopy (FLIM)

Cells were grown overnight in Synthetic Complete medium to OD_600_ = 0.05 to 0.1, concentrated by spinning and incubated for 1min with 2ng/mL of the Flipper-TR probe. FLIM imaging was performed using a Nikon Eclipse Ti A1R microscope equipped with a time-correlated single-photon counting module from PicoQuant58. Excitation was performed using a pulsed 485 nm laser (PicoQuant, LDH-D-C-485) operating at 20 MHz, and the emission signal was collected through a 600/50 nm bandpass filter using a gated PMA hybrid 40 detector and a TimeHarp 260 PICO board (PicoQuant).

### Image processing and quantification

The acquired image series were background subtracted and corrected for general photobleaching. Final image processing and analysis were done using ImageJ.

For FLIM analysis (Fig. 4A, Fig S3A-B, Fig 5A), we used the SymPhoTime 64 software (PicoQuant) to fit the data according to a 2-exponential reconvolution model and calculate the lifetime of the Flipper-TR probe.

### Statistics and reproducibility

The sample sizes and statistical tests were selected based on previous studies with similar methodologies. All experiments were repeated at least three times, giving similar results. The results of independent experiments are presented as mean values; error bars represent the SD, or the propagated error when the value of each experiment was itself calculated as a mean of individual cells. Statistical significance was tested using the two-tailed Student’s t-test.

## Data Availability

All data that support the findings of this study are available from the corresponding authors upon reasonable request. Correspondence and requests for materials should be addressed to Robbie Loewith: or Aurélien Roux

## Contributions

MR performed the experiments and wrote the first draft of the manuscript. MM synthesized the Flipper-TR probe under the direction and support of SM. MR, RL and AR designed the experiments and edited the manuscript.

## Acknowledgements

SM, RL and AR acknowledge generous financial support from the Canton of Geneva, the Swiss National Science Foundation and the NCCR in Chemical Biology.

---

**Figure S1:**
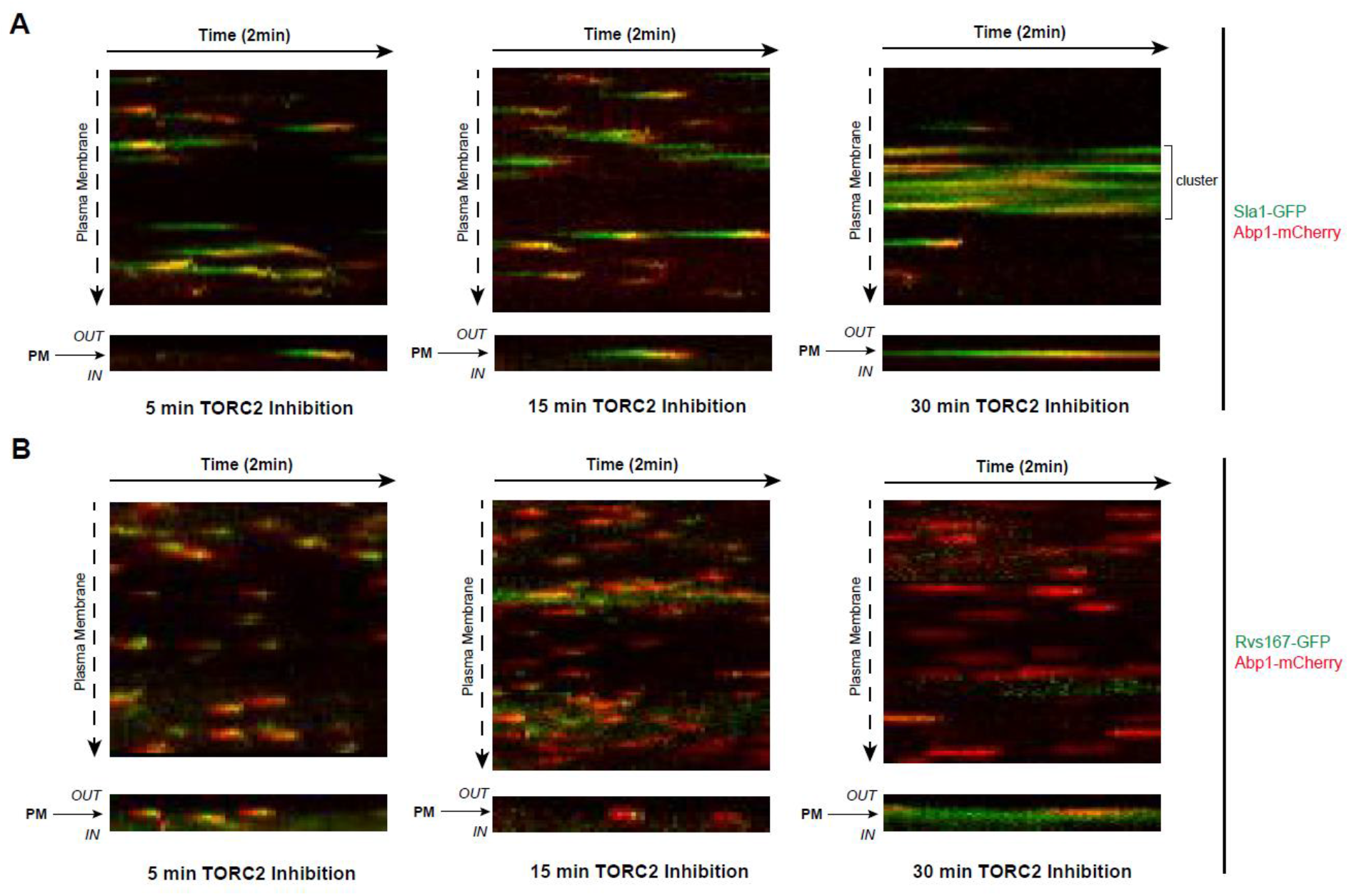
TORC2 inhibition effects on endocytosis are progressive. Kymographs of *TOR1-1 AVO3Δ^CT^* cells tagged with Abp1-mCherry and either Sla1-GFP **(A)** or Rvs167-GFP **(B)**, recorded after the indicated time of Rapamycin treatment. The kymographs were recorded either along (top) or across (bottom) the plasma membrane.

**Figure S2:**
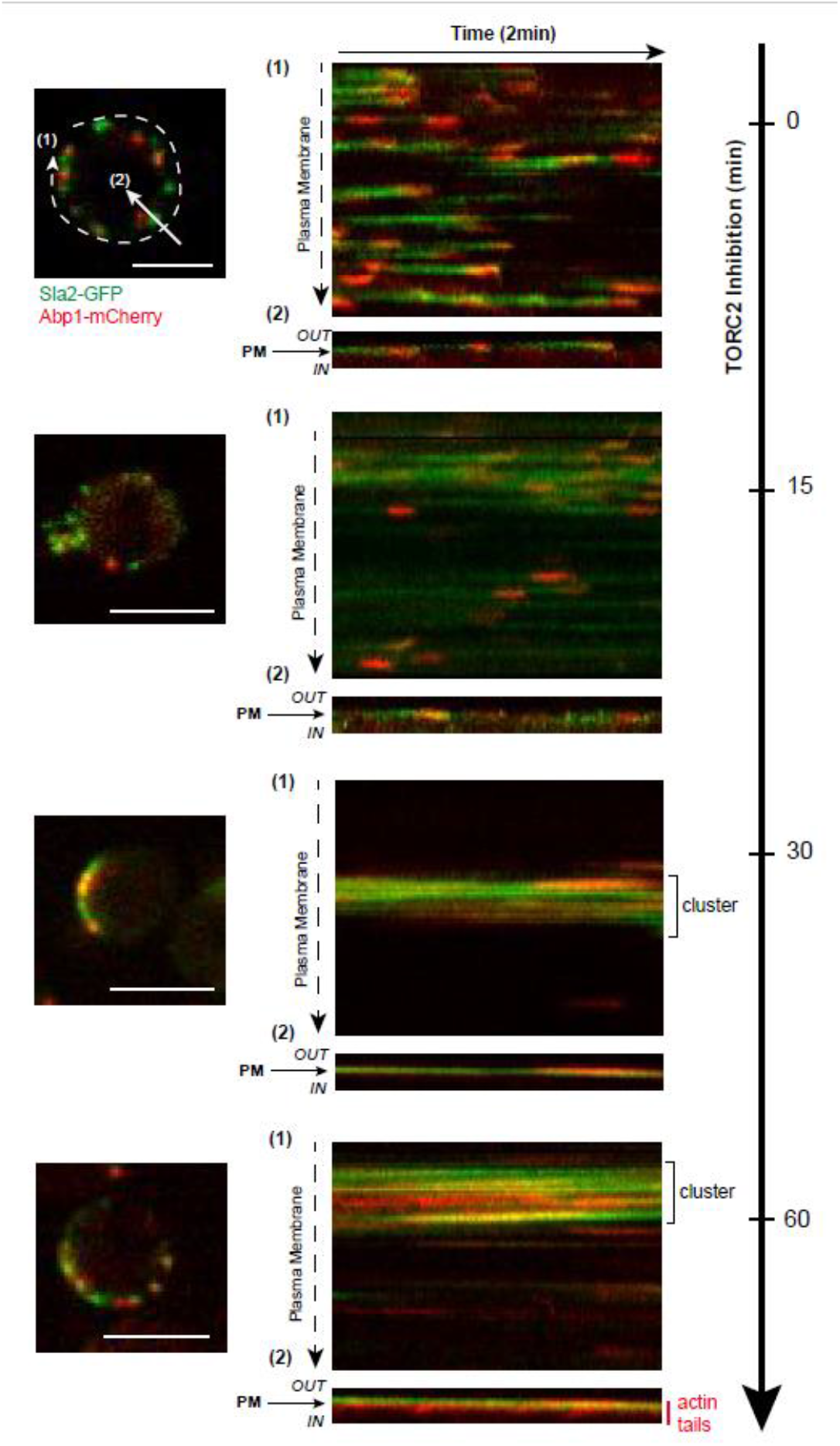
Sla2 does not colocalize with actin tails upon TORC2 inhibition. *(Left)* Merged images of Sla2-GFP and Abp1-mCherry in *TOR1-1 AVO3Δ^CT^* cells upon TORC2 inhibition by Rapamycin treatment. *(Right)* Sla2-GFP and Abp1-mCherry kymographs of the indicated cells, after the indicated time of TORC2 inhibition. Kymographs were recorded either along (1) or across (2) the Plasma Membrane, as explained on the left pictures. All scale bars, 5μm.

**Figure S3:**
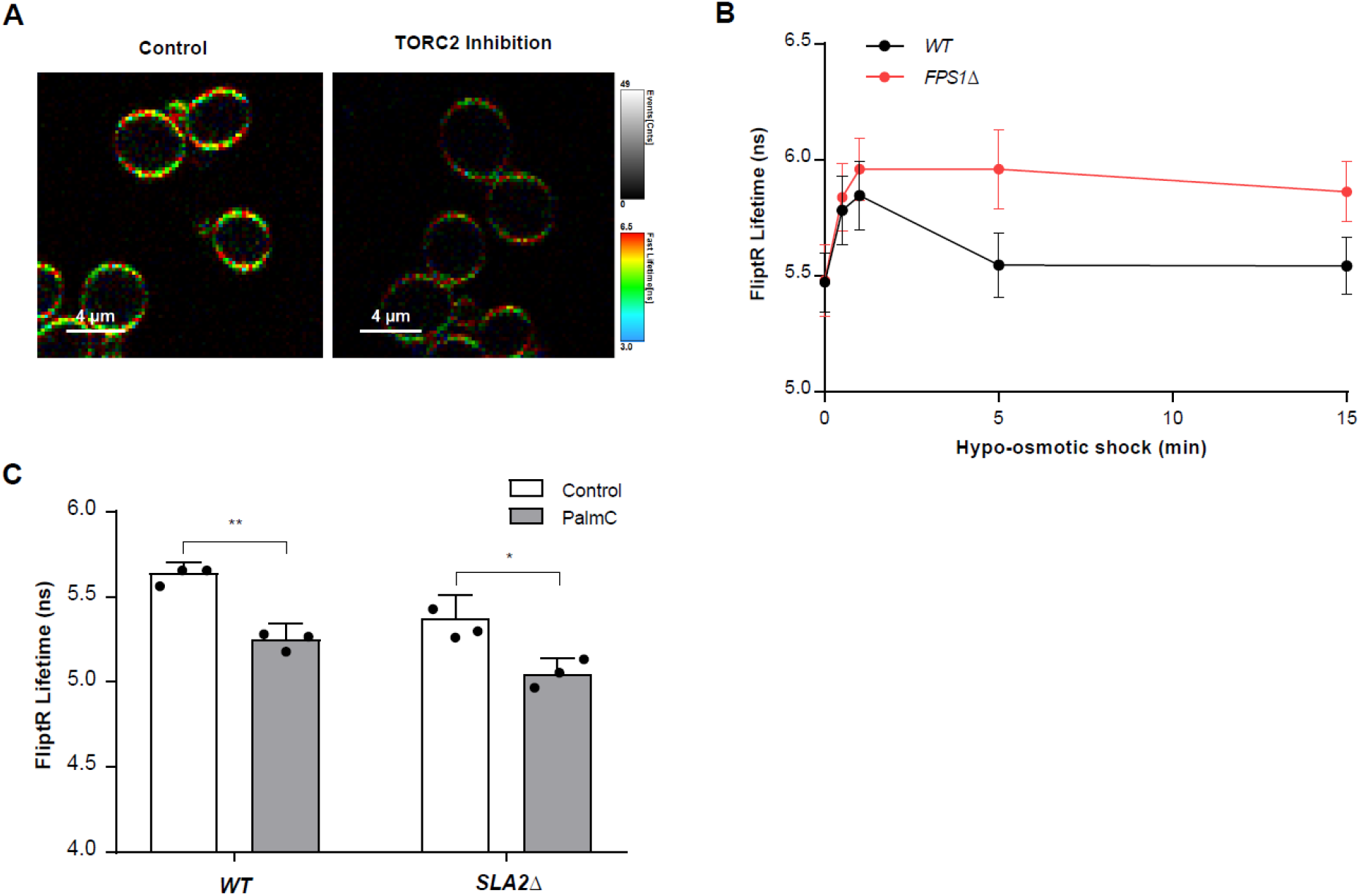
Osmotic shocks and PalmC treatment affect Plasma Membrane tension in yeast. **(A)** Color-coded FLIM images of *TOR1-1 AVO3Δ^CT^* cells labelled with FliptR, before and after 1h Rapamycin treatment. Scale bars, 4μM. **(B)** Evolution of PM tension, monitored through the lifetime of the FliptR probe measured by FLIM, upon hypo-osmotic shock in *WT* or *FPS1Δ* cells. Error bars represent the propagated error of mean values calculated from three independent experiments (with n>10 cells). **(C)** PM tension, monitored through the lifetime of the FliptR probe measured by FLIM, before and after PalmC treatment in *WT* or *SLA2Δ* cells. Error bars represent the propagated error of mean values calculated from three independent experiments with n>10 cells (** p<0.01; * p<0.05).

**Supplementary Table 1:**
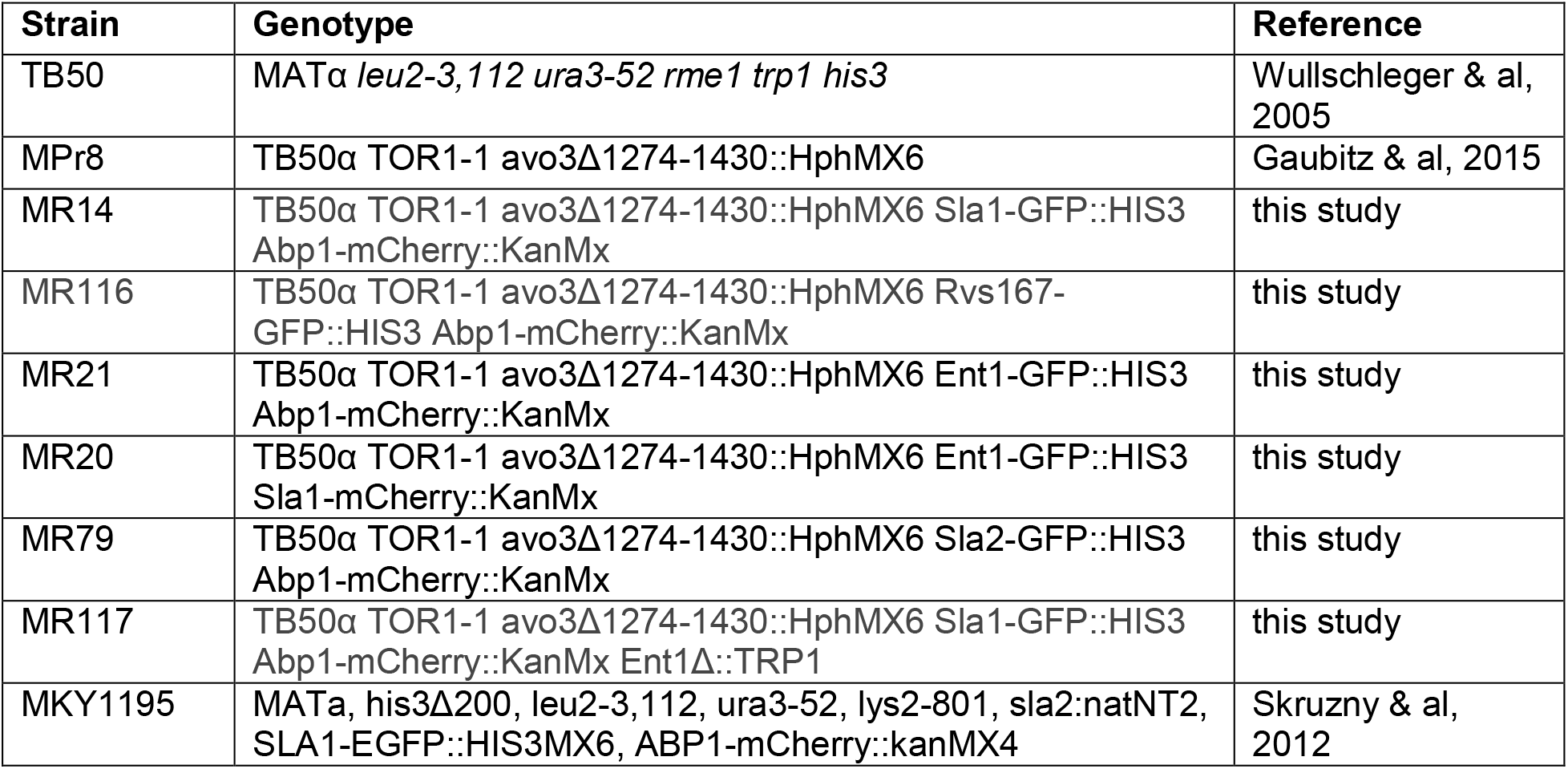
Strains used in this study

**Supplementary Table 2:** Plasmids used in this study

| Plasmid | Description | Reference |
| --- | --- | --- |
| p525 | pFA6a-mCherry-kan | this study |
| p16 | pFA6a-GFP(S65T)-HIS3 | Longtine M.S. & al(1989), Yeast 14, 953-961 |
| p12 | pFA6a-TRP1 | Longtine M.S. & al(1989), Yeast 14, 953-961 |

**Supplementary Table 3:**
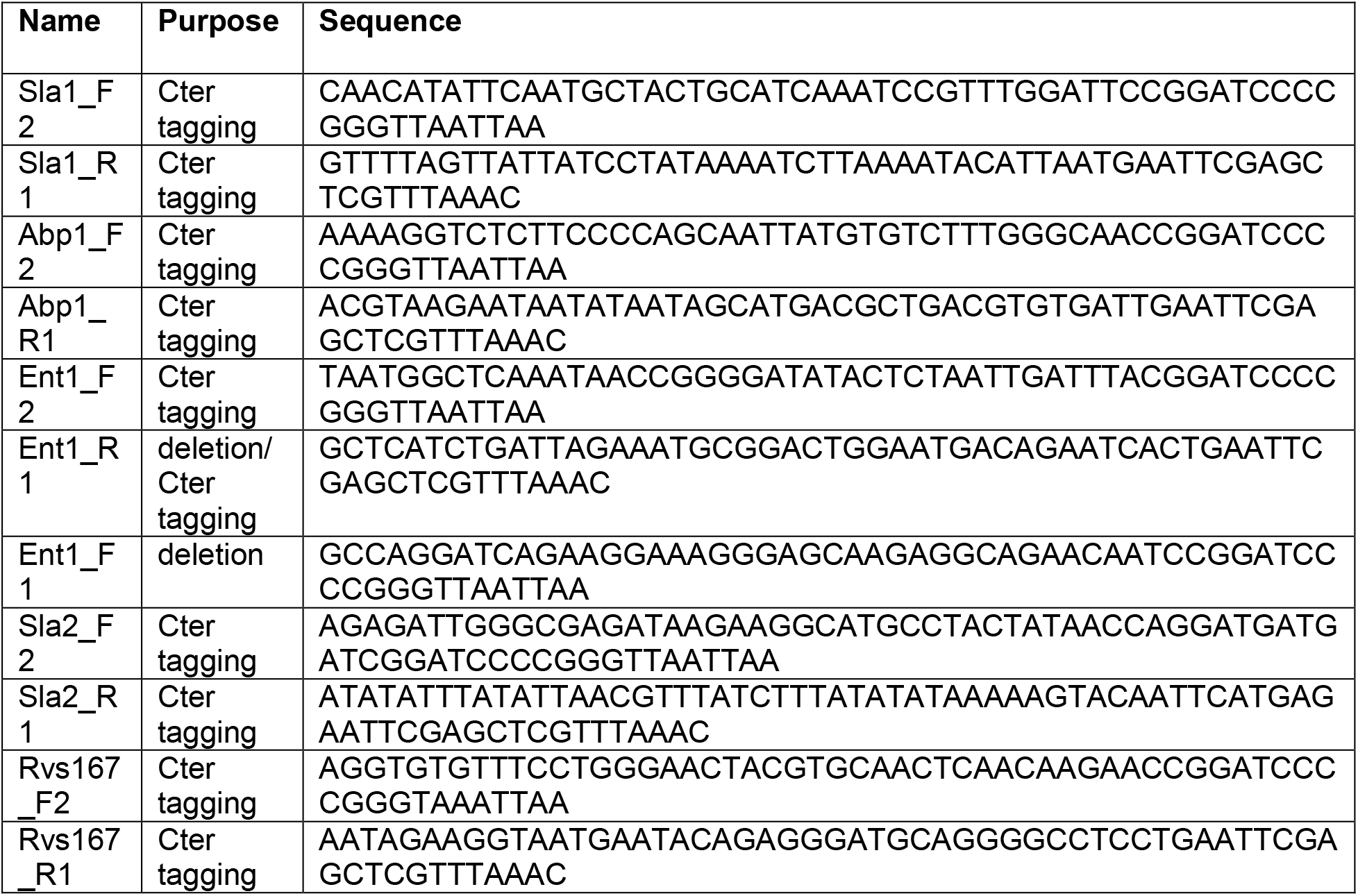
Primers used in this study

## References

Aghamohammadzadeh, S., and Ayscough, K.R. (2009). Differential requirements for actin during yeast and mammalian endocytosis. Nat Cell Biol 11, 1039–1042.

Apodaca, G. (2002). Modulation of membrane traffic by mechanical stimuli. Am J Physiol Renal Physiol 282, F179–190.

Aronova, S., Wedaman, K., Aronov, P.A., Fontes, K., Ramos, K., Hammock, B.D., and Powers, T. (2008). Regulation of ceramide biosynthesis by TOR complex 2. Cell Metab 7, 148–158.

Basu, R., Munteanu, E.L., and Chang, F. (2014). Role of turgor pressure in endocytosis in fission yeast. Mol Biol Cell 25, 679–687.

Beese, S.E., Negishi, T., and Levin, D.E. (2009). Identification of positive regulators of the yeast fps1 glycerol channel. PLoS Genet 5, e1000738.

Berchtold, D., Piccolis, M., Chiaruttini, N., Riezman, I., Riezman, H., Roux, A., Walther, T.C., and Loewith R. (2012). Plasma membrane stress induces relocalization of Slm proteins and activation of TORC2 to promote sphingolipid synthesis. Nat Cell Biol 14, 542–547.

Boulant, S., Kural, C., Zeeh, J.C., Ubelmann, F., and Kirchhausen, T. (2011). Actin dynamics counteract membrane tension during clathrin-mediated endocytosis. Nat Cell Biol 13, 1124–1131.

Bourgoint, C., Rispal, D., Berti, M., Filipuzzi, I., Helliwell, S.B., Prouteau, M., and Loewith, R. (2018). Target of rapamycin complex 2-dependent phosphorylation of the coat protein Pan1 by Akl1 controls endocytosis dynamics in Saccharomyces cerevisiae. J Biol Chem 293, 12043–12053.

Brady, R.J., Damer, C.K., Heuser, J.E., and O’Halloran, T.J. (2010). Regulation of Hip1r by epsin controls the temporal and spatial coupling of actin filaments to clathrin-coated pits. J Cell Sci 123, 3652–3661.

Breslow, D.K., Collins, S.R., Bodenmiller, B., Aebersold, R., Simons, K., Shevchenko, A., Ejsing, C.S., and Weissman, J.S. (2010). Orm family proteins mediate sphingolipid homeostasis. Nature 463, 1048–1053.

Carroll, S.Y., Stimpson, H.E., Weinberg, J., Toret, C.P., Sun, Y., and Drubin, D.G. (2012). Analysis of yeast endocytic site formation and maturation through a regulatory transition point. Mol Biol Cell 23, 657–668.

Colom, A., Derivery, E., Soleimanpour, S., Tomba, C., Molin, M.D., Sakai, N., Gonzalez-Gaitan, M., Matile, S., and Roux, A. (2018). A fluorescent membrane tension probe. Nat Chem 10, 1118–1125.

Dai, J., and Sheetz, M.P. (1995). Mechanical properties of neuronal growth cone membranes studied by tether formation with laser optical tweezers. Biophys J 68, 988–996.

deHart, A.K., Schnell, J.D., Allen, D.A., and Hicke, L. (2002). The conserved Pkh-Ypk kinase cascade is required for endocytosis in yeast. J Cell Biol 156, 241–248.

deHart, A.K., Schnell, J.D., Allen, D.A., Tsai, J.Y., and Hicke, L. (2003). Receptor internalization in yeast requires the Tor2-Rho1 signaling pathway. Mol Biol Cell 14, 4676–4684.

Diz-Munoz, A., Thurley, K., Chintamen, S., Altschuler, S.J., Wu, L.F., Fletcher, D.A., and Weiner, O.D. (2016). Membrane Tension Acts Through PLD2 and mTORC2 to Limit Actin Network Assembly During Neutrophil Migration. PLoS Biol 14, e1002474.

Dmitrieff, S., and Nedelec, F. (2015). Membrane Mechanics of Endocytosis in Cells with Turgor. PLoS Comput Biol 11, e1004538.

Engqvist-Goldstein, A.E., Zhang, C.X., Carreno, S., Barroso, C., Heuser, J.E., and Drubin, D.G. (2004). RNAi-mediated Hip1R silencing results in stable association between the endocytic machinery and the actin assembly machinery. Mol Biol Cell 15, 1666–1679.

Fernandez-Sanchez, M.E., Brunet, T., Roper, J.C., and Farge, E. (2015). Mechanotransduction’s impact on animal development, evolution, and tumorigenesis. Annu Rev Cell Dev Biol 31, 373–397.

Gaubitz, C., Oliveira, T.M., Prouteau, M., Leitner, A., Karuppasamy, M., Konstantinidou, G., Rispal, D., Eltschinger, S., Robinson, G.C., Thore, S., et al. (2015). Molecular Basis of the Rapamycin Insensitivity of Target Of Rapamycin Complex 2. Mol Cell 58, 977–988.

Gauthier, N.C., Masters, T.A., and Sheetz, M.P. (2012). Mechanical feedback between membrane tension and dynamics. Trends Cell Biol 22, 527–535.

Godlee, C., and Kaksonen, M. (2013). Review series: From uncertain beginnings: initiation mechanisms of clathrin-mediated endocytosis. J Cell Biol 203, 717–725.

Gottlieb, T.A., Ivanov, I.E., Adesnik, M., and Sabatini, D.D. (1993). Actin microfilaments play a critical role in endocytosis at the apical but not the basolateral surface of polarized epithelial cells. J Cell Biol 120, 695–710.

Kaksonen, M., and Roux, A. (2018). Mechanisms of clathrin-mediated endocytosis. Nat Rev Mol Cell Biol 19, 313–326.

Kaksonen, M., Sun, Y., and Drubin, D.G. (2003). A pathway for association of receptors, adaptors, and actin during endocytic internalization. Cell 115, 475–487.

Kaksonen, M., Toret, C.P., and Drubin, D.G. (2005). A modular design for the clathrin-and actin-mediated endocytosis machinery. Cell 123, 305–320.

Kaksonen, M., Toret, C.P., and Drubin, D.G. (2006). Harnessing actin dynamics for clathrin-mediated endocytosis. Nat Rev Mol Cell Biol 7, 404–414.

Kukulski, W., Schorb, M., Kaksonen, M., and Briggs, J.A. (2012). Plasma membrane reshaping during endocytosis is revealed by time-resolved electron tomography. Cell 150, 508–520.

Morlot, S., Galli, V., Klein, M., Chiaruttini, N., Manzi, J., Humbert, F., Dinis, L., Lenz, M., Cappello, G., and Roux, A. (2012). Membrane shape at the edge of the dynamin helix sets location and duration of the fission reaction. Cell 151, 619–629.

Morris, C.E., and Homann, U. (2001). Cell surface area regulation and membrane tension. J Membr Biol 179, 79–102.

Muir, A., Ramachandran, S., Roelants, F.M., Timmons, G., and Thorner, J. (2014). TORC2-dependent protein kinase Ypk1 phosphorylates ceramide synthase to stimulate synthesis of complex sphingolipids. Elife 3.

Picco, A., Kukulski, W., Manenschijn, H.E., Specht, T., Briggs, J.A.G., and Kaksonen, M. (2018). The contributions of the actin machinery to endocytic membrane bending and vesicle formation. Mol Biol Cell 29, 1346–1358.

Riggi, M., Niewola-Staszkowska, K., Chiaruttini, N., Colom, A., Kusmider, B., Mercier, V., Soleimanpour, S., Stahl, M., Matile, S., Roux, A., et al. (2018). Decrease in plasma membrane tension triggers PtdIns(4,5)P2 phase separation to inactivate TORC2. Nat Cell Biol 20, 1043–1051.

Rispal, D., Eltschinger, S., Stahl, M., Vaga, S., Bodenmiller, B., Abraham, Y., Filipuzzi, I., Movva, N.R., Aebersold, R., Helliwell, S.B., et al. (2015). Target of Rapamycin Complex 2 Regulates Actin Polarization and Endocytosis via Multiple Pathways. J Biol Chem 290, 14963–14978.

Roelants, F.M., Breslow, D.K., Muir, A., Weissman, J.S., and Thorner, J. (2011). Protein kinase Ypk1 phosphorylates regulatory proteins Orm1 and Orm2 to control sphingolipid homeostasis in Saccharomyces cerevisiae. Proc Natl Acad Sci U S A 108, 19222–19227.

Roelants, F.M., Leskoske, K.L., Pedersen, R.T., Muir, A., Liu, J.M., Finnigan, G.C., and Thorner, J. (2017). TOR Complex 2-Regulated Protein Kinase Fpk1 Stimulates Endocytosis via Inhibition of Ark1/Prk1-Related Protein Kinase Akl1 in Saccharomyces cerevisiae. Mol Cell Biol 37.

Saleem, M., Morlot, S., Hohendahl, A., Manzi, J., Lenz, M., and Roux, A. (2015). A balance between membrane elasticity and polymerization energy sets the shape of spherical clathrin coats. Nat Commun 6, 6249.

Simunovic, M., Voth, G.A., Callan-Jones, A., and Bassereau, P. (2015). When Physics Takes Over: BAR Proteins and Membrane Curvature. Trends Cell Biol 25, 780–792.

Skruzny, M., Brach, T., Ciuffa, R., Rybina, S., Wachsmuth, M., and Kaksonen, M. (2012). Molecular basis for coupling the plasma membrane to the actin cytoskeleton during clathrin-mediated endocytosis. Proc Natl Acad Sci U S A 109, E2533–2542.

Skruzny, M., Desfosses, A., Prinz, S., Dodonova, S.O., Gieras, A., Uetrecht, C., Jakobi, A.J., Abella, M., Hagen, W.J., Schulz, J., et al. (2015). An organized co-assembly of clathrin adaptors is essential for endocytosis. Dev Cell 33, 150–162.

Watson, H.A., Cope, M.J., Groen, A.C., Drubin, D.G., and Wendland, B. (2001). In vivo role for actin-regulating kinases in endocytosis and yeast epsin phosphorylation. Mol Biol Cell 12, 3668–3679.

Wendland, B., Steece, K.E., and Emr, S.D. (1999). Yeast epsins contain an essential N-terminal ENTH domain, bind clathrin and are required for endocytosis. EMBO J 18, 4383–4393.

Youn, J.Y., Friesen, H., Kishimoto, T., Henne, W.M., Kurat, C.F., Ye, W., Ceccarelli, D.F., Sicheri, F., Kohlwein, S.D., McMahon, H.T., et al. (2010). Dissecting BAR domain function in the yeast Amphiphysins Rvs161 and Rvs167 during endocytosis. Mol Biol Cell 21, 3054–3069.

